# Elimination of HSV-2 infected cells is mediated predominantly by paracrine effects of tissue-resident T cell derived cytokines

**DOI:** 10.1101/610634

**Authors:** Pavitra Roychoudhury, David A Swan, Elizabeth Duke, Lawrence Corey, Jia Zhu, Veronica Davé, Laura Richert Spuhler, Jennifer M. Lund, Martin Prlic, Joshua T. Schiffer

## Abstract

The mechanisms underlying rapid elimination of herpes simplex virus-2 (HSV-2) in the human genital tract despite low tissue-resident CD8+ T-cell density (T_RM_) are unknown. We analyzed shedding episodes during chronic HSV-2 infection: viral clearance always occurred within 24 hours of detection even if viral load exceeded 10^7^ HSV DNA copies; surges in granzyme B and interferon-*γ* occurred within the early hours after reactivation. We next developed a mathematical model of an HSV-2 genital ulcer to integrate mechanistic observations of T_RM_ *in situ* proliferation, trafficking, cytolytic effects and cytokine alarm signaling from murine studies with viral kinetics, histopathology and lesion size data from humans. A sufficiently high density of HSV-2 specific T_RM_ predicted rapid contact-mediated elimination of infected cells. At lower T_RM_ densities, T_RM_ must initiate a rapidly diffusing, polyfunctional cytokine response in order to eliminate of a majority of infected cells and eradicate briskly spreading HSV-2 infection.

**One Sentence Summary:** Control of herpes simplex virus-2 is primarily mediated by rapidly diffusing cytokines secreted by tissue-resident T cells.

## Introduction

Tissue resident memory CD8+ T cells (T_RM_) rapidly identify and control recurrent human viral infections within peripheral sites (*1-11*). T_RM_ lodge preferentially at prior sites of pathogen replication without recirculating in blood, and are differentiated from circulating memory CD8+ T cells based on micro-anatomic localization, cell-surface markers, and transcriptional profile(*5, 12-15*). During HSV-2 infection in humans, cells with T_RM_ characteristics cluster at the dermal-epidermal junction where neuron endings release HSV-2 into the mucosa (*9, 10, 16*). The heterogeneous spatial aggregation of T_RM_ in genital skin explains the considerable variability of HSV-2 severity, including episodes which are contained after a few hours of viral replication and those that persist for many days and cause symptomatic lesions (*17-21*).

Recent murine studies employing intravital imaging describe the extraordinary range of T_RM_ control mechanisms *in situ*. In the absence of antigen, T_RM_ efficiently survey previously infected tissues to recognize and eliminate infected cells during re-infection (*1, 22*). During this immunosurveillance phase, T_RM_ express dendritic-like arms to contact a large number of potentially infected cells (*1, 22*), and remain in an activated state despite the absence of antigen-driven stimulus via the T-cell receptor (*8*). Upon recognition of an infected cell, T_RM_ motility is restricted and abundant local proliferation ensues (*22*). T_RM_ then mediate efficient, contact-mediated apoptosis of infected cells by releasing perforin and granzyme B (*23-25*).

Upon activation, T_RM_ induce a broad antiviral program consisting of cytokine release and resultant generalized local inflammation (*1, 8*). Cytokines, notably interferon-*γ* (IFN-*γ*), induce clearance of infected cells (*26, 27*), and render surrounding cells resistant to infection, even with unrelated pathogens (*1*). Chemokines such as CXCL9 and 10 may draw local immune cells to infected sites (*11, 28*). Cytokines and chemokines diffuse rapidly, are efficiently internalized by their target cells (*29*) and exert paracrine effects upon cells that are not in direct contact with T_RM_.

Analysis of human biopsy specimens of HSV-2 infected tissues reveals high-density clusters of T_RM_ in certain micro-regions, with low densities of T_RM_ in others (*20*). Yet, a majority of HSV-2 shedding episodes are eliminated within 6-24 hours (*19, 21, 30*). Because individual HSV-2 ulcers are small, this rapid peak in viral load is more likely due to an intense local immune response rather than target cell limitation (*19, 30-32*).

It is unknown how low numbers of T_RM_ orchestrate elimination of infected cells so efficiently. Different mechanisms may mediate elimination of a few infected cells over hours during mild episodes, versus thousands of infected cells over days during more severe episodes. Here, we construct an agent-based mathematical model to recapitulate the spatiotemporal dynamics underlying the viral spread, cytokine expansion and T_RM_-mediated containment of HSV-2 in genital tissues.

## Results

### Intense immune pressure within 24 hours of HSV-2 detection

A characteristic of HSV-2 shedding episodes in chronically infected persons is rapid viral expansion, followed by a slightly more protracted clearance phase (*21, 33, 34*). While most episodes share this core feature, they vary in severity. Many asymptomatic episodes are cleared in 6-12 hours and peak at <10^4^ HSV DNA copies. Other episodes exceed 10^7^ HSV DNA copies and are associated with visible lesions and shedding over days to weeks (*30, 35, 36*). In episodes longer than four days, longevity is driven by a rebound of HSV after an initial phase of clearance, possibly due to seeding of new ulcers in immunologically distinct micro-environments (*19*).

To approximate viral dynamics within single infection micro-environments, we re-analyzed 83 episodes (655 total swabs from 20 subjects) from a dataset in which study participants performed swabs every 6 hours over 60 days. We binned episodes by duration into short (<1 day), medium (1–2 days), and long (>2 days) (*30*), and identified brisk viral expansion with an early peak in all episodes, even those with high viral loads **(Fig. 1a)**. There was a wide distribution of peak viral loads **(Fig. 1b)** ranging from 2.2 to 7.9 log10 HSV DNA copies. Episodes were of variable duration, but elimination usually occurred in less than 48 hours, with a minority lasting >100 hours **(Fig. 1c)**. Median time to the first viral load peak was 7 hours; all episodes reached their first peak within 24 hours **(Fig. 1d)**. Moderate correlations were noted between time to first peak and peak viral load **(Fig. 1e)**, and time to first peak and duration **(Fig. 1f)**.

**Fig 1.**
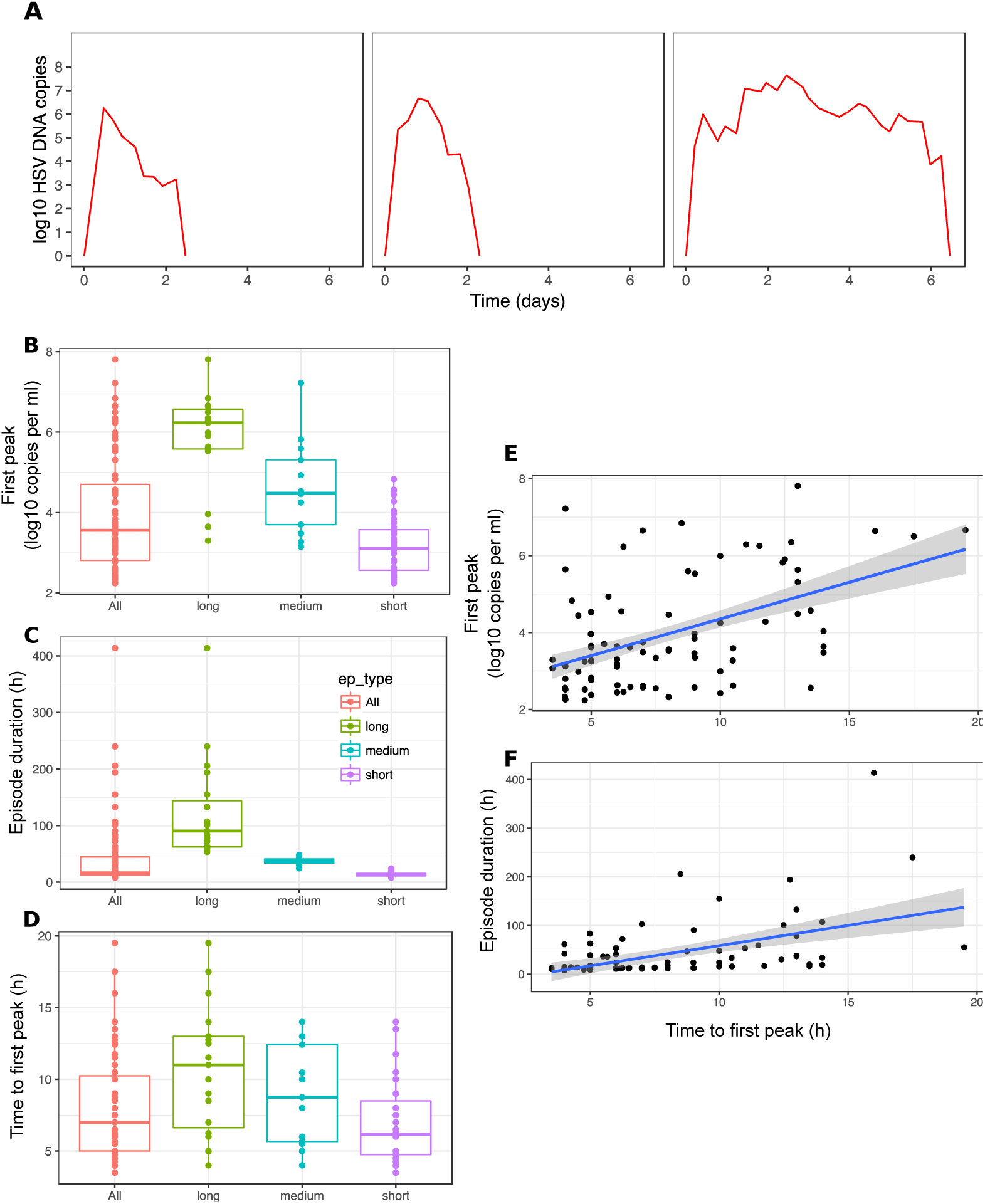
Early and intense immunological pressure against HSV-2 replication within genital micro-environments. **(A)** Three severe shedding episodes with typical characteristics including rapid time to first peak viral load and rapid viral clearance. **(B-D)** Analysis of 83 shedding episodes derived from every 6-hour genital sampling. Boxplots with interquartile range or IQR (box) and 1.5x the IQR (whiskers). Categorization according to duration: short <24 hours (n=51), medium 24-48 hours (n=13), and long >48 hours (n=19). **(B)** Variable peak viral loads and **(C)** durations across episodes with most episodes eliminated in fewer than 48 hours. **(D)** Uniform occurrence of first peak viral load within the first 24 hours of HSV-2 detection. **(E)** Correlation of time to peak viral load with peak episode viral load (Spearman’s ρ=0.434, p<0.001). **(F)** Correlation of time to peak viral load with episode duration (Spearman’s ρ=0.534, p<0.001).

These results confirm that massive HSV-2 replication is a defining early feature of shedding episodes. However, immunologic responses predominate within 20 hours of viral detection, and time of the initial peak predicts severity. Whether an episode progresses to high viral loads and days of shedding or rapid containment in the first several hours may depend on the intensity of the immune response during the early hours of reactivation.

### Heterogeneous dispersion of tissue-resident CD8+ T cells (T_RM_) in HSV-2 infected genital tissues

HSV-2 reactivates in specific micro-environments throughout the human genital tract because of random travelling of virions along highly arborized branches of peripheral neurons (*37*). We observed a fixed spatial meta-structure of CD8+ T-cell density in infected tissue, defined by dense clusters in some regions and low numbers in others (*20*). To characterize tissue micro-environments, we summarized T_RM_ densities observed in 19 genital biopsy specimens **(Fig. 2a)** performed at 2 and 8 weeks following lesion healing (We refer to all observed CD8+ T cells as T_RM_ though it is possible that a minority are not truly resident). We divided each image into regions with 100 epidermal cells per region **(Fig. 2b-c)** and calculated the ratio of T_RM_ to epidermal cells (*in situ* effector:target or E:T ratio). We then created a rank-order distribution of all observed ratios **(Fig. 2d)**. The median E:T ratio in the 1008 micro-regions was 0.02 (IQR 0.07, range 0-2). T_RM_ were absent in 26.2% of the micro-regions (E:T=0). Median E:T ratios for the 19 specimens ranged from 0.01 to 0.24 (IQR of medians = 0.02). Exponential slopes of rank order abundance curves (median −0.121, IQR 0.048, range: −0.222 to −0.045) were similar across specimens **(Fig. 2d)**, implying an equivalent gradient between high and low T_RM_ density regions **(Fig. 2a, c)**. Specimens with a higher proportion of micro-regions with E:T=0 had lower overall E:T ratios **(Fig. 2e)**.

**Fig 2.**
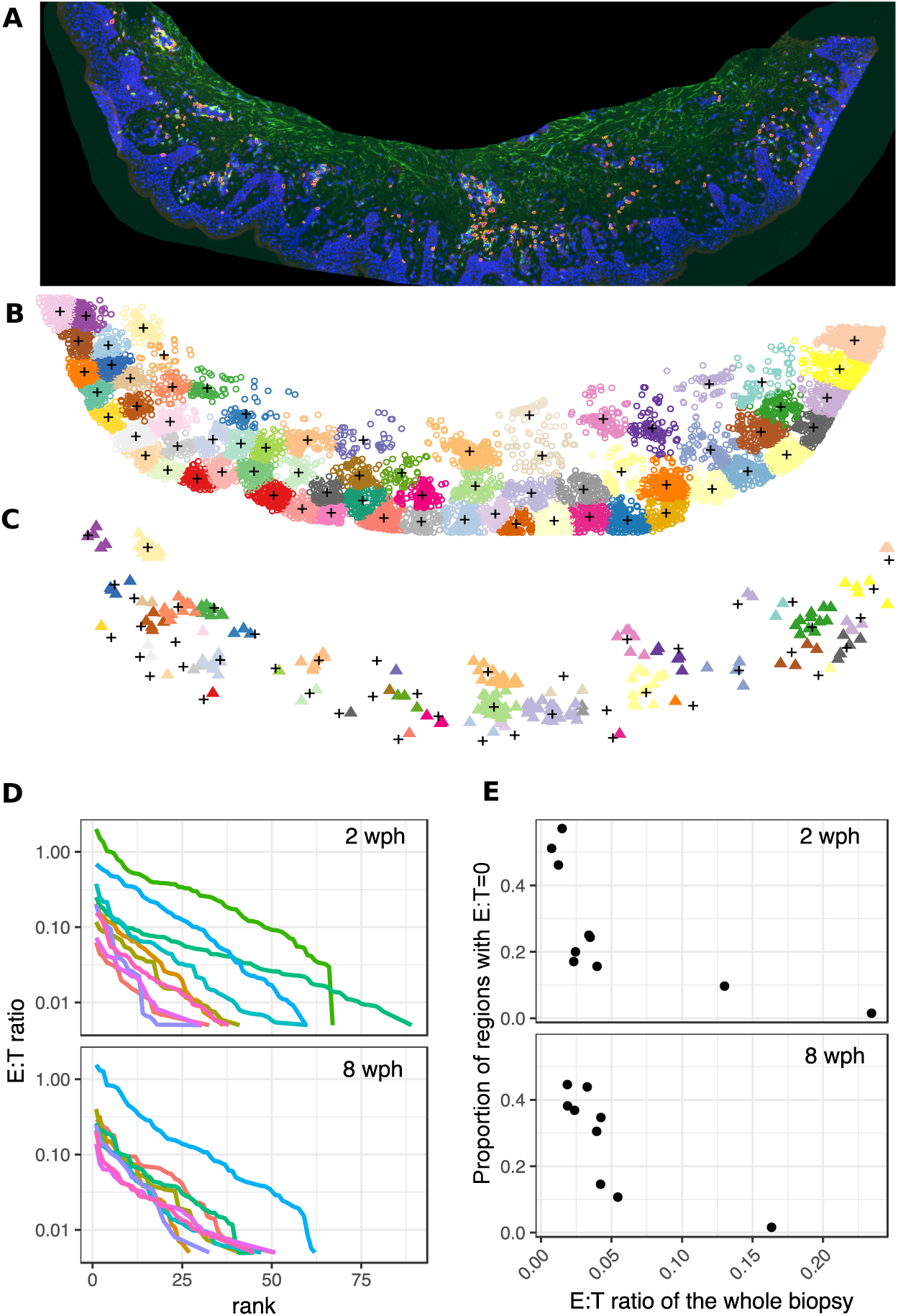
Heterogeneous dispersion of CD8+ T cells within HSV-2 infected genital biopsy specimens obtained after lesion healing. Analysis of 19 genital biopsy specimens from 10 participants 2 weeks (n=10) or 8 weeks (n=9) after lesion healing. **(A)** Three-color RGB image with cell nuclei (blue), CD4+ T cells (green), and CD8+ T cells (red) of a 2-week specimen. **(B)** 100-epidermal cell micro-regions designated according to clusters shown as separate colors; “+” = cluster center. **(C)** CD8+ T cells (triangles) within each micro-region. **(D)** Rank order distributions of CD8+ T cell : epithelial (E:T) ratios within micro-regions; lines = single specimens; wide ranges of E:T ratios within each specimen with similar rank order slopes across specimens. **(E)** High variability of whole specimen E:T ratios (x-axis) across 2- and 8-week specimens; inverse correlation of whole specimen E:T ratios with the proportion of 100 keratinocyte regions containing no CD8+ T cells (y-axis).

### A mathematical model of HSV-2 replication and spread within a single ulcer

We next sought to explain how infected cells are eliminated quickly, given the low T_RM_ densities commonly observed in infected samples. To approximate spatial conditions of a single infection micro-environment, we developed a two-dimensional stochastic, agent-based mathematical model. Our goal was to precisely link spatial heterogeneity of T_RM_ density **(Fig. 2)** with observed heterogeneity in shedding episode kinetics **(Fig. 1)**. We developed the model in an iterative fashion to limit required parameter fitting at each step.

First, we simulated HSV-2 replication and spread assuming no T_RM_ immunity (**Fig. 3a**, **Materials and Methods** for details and **Table 1** for model parameters), to recapitulate early HSV-2 shedding episode kinetics. The agent-based model assumes that HSV-2 replication occurs in a two-stage process within an infected cell. Newly infected cells enter a pre-productive phase, during which immediate-early and early viral proteins are expressed but HSV DNA replication has yet to occur (*38*). After several hours, HSV DNA polymerase activity initiates, leading to production of infectious viral particles for ∼20 hours until cell lysis and death **(Fig. 3a)**.

**Table 1.** Model parameters for cell and virus compartments in the absence of immunity

| Parameter | Unit | Value | Source |
| --- | --- | --- | --- |
| Uninfected cell div rate<sup>1</sup> ($\alpha$) | day<sup>-1</sup> | 0.077 | (61) |
| Uninfected cell death rate<sup>2</sup> ($\delta_S$) | day<sup>-1</sup> | 0 | |
| Infected cell death rate ($\delta_I$) | day<sup>-1</sup> | 1.25 | (60) |
| Lag in virus production<sup>3</sup> ($\varepsilon$) | cell<sup>-1</sup> day<sup>-1</sup> | 8 | (62) |
| Infectivity ($\beta$) | virus<sup>-1</sup> cell<sup>-1</sup> day<sup>-1</sup> | 0.1 | Fitted (initial model) |
| Virus spread rate<sup>4</sup> | sites day<sup>-1</sup> | 54 | Fitted (final model) |
| Virus production rate<br>($\mu$, mean ± SD) | cell<sup>-1</sup> day<sup>-1</sup> | $3.16 \pm 1.00 \times 10^4$ | (60) |
| Free virus decay rate | site<sup>-1</sup> day<sup>-1</sup> | 8.8 | (19) |
| Spatial scale | $\mu\text{m site}^{-1}$ | 50 | (63, 64) |

1 Rate at which a vacant site becomes occupied by a susceptible cell
2 We assumed that cell death due to infection greatly exceeds uninfected cell death during lesion formation.
3 This rate represents the time from infection to first released virus
4 Rate at which infectious virus spreads across the grid

**Fig 3.**
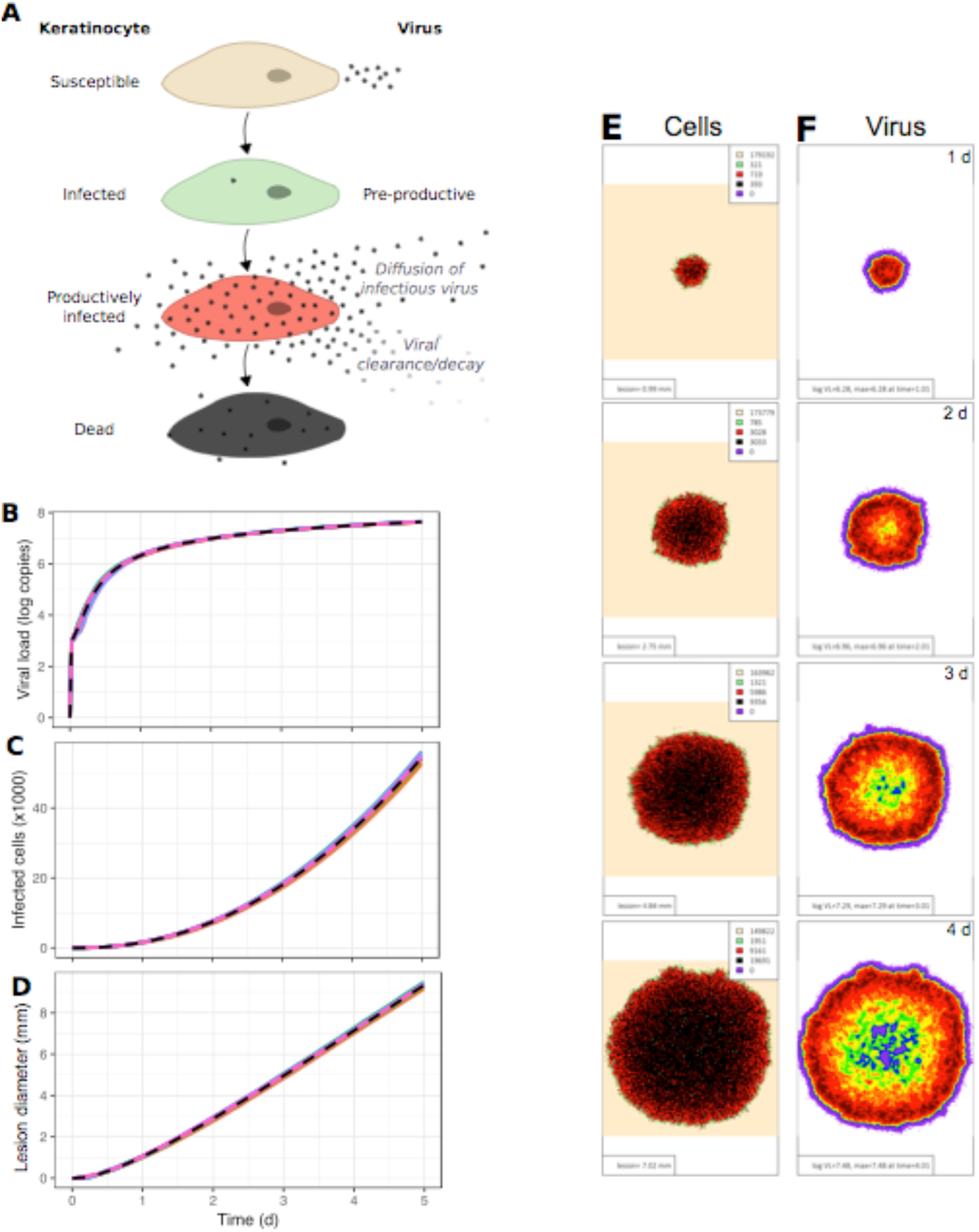
Continual spatial expansion of HSV-2 infection in mathematical model simulations without T_RM_. **(A)** Mathematical model of HSV-2 replication and spread; infected cell progression through pre-productive (green), and productive phases (red) during which viral replication occurs and virus diffuses to surrounding regions. **(B)** HSV-2 viral load trajectory versus time for 10 simulated episodes with asymptotic behavior and no viral elimination (dotted line = median). **(C)** Number of infected cells and **(D)** ulcer diameter versus time with continual expansion in the absence of a T_RM_ response in the 10 simulated episodes. **(E-F)** Spatial approximation of infection spread in 5-day simulations which correspond to **Movie S1**; one-day time steps per row. **(E)** An expanding ulcer with a leading edge of pre-productively infected cells (green), an inner ring of productively infected cells (red) and a core of dead cells (black). **(F)** Viral diffusion extending beyond the range of infected cells with highest local viral loads noted over regions with active HSV-2 replication; viral loads (HSV DNA copies) per cellular region: purple=10^2^-10^2^.^25^, blue=10^2^.^25^-10^2^.^5^, green=10^2^.^5^-10^2^.^75^, yellow=10^2^.^75^-10^3^, orange=10^3^-10^3^.^25^, red=10^3^.^25^-10^3^.^5^, dark red>10^3^.^5^.

Human HSV-2 infection occurs within nucleated epithelial cells in genital skin or mucosa. The three-dimensional anatomy of this tissue space comprises ∼2-10 stacked epidermal cells between the surface of the skin and the dermal-epidermal junction where HSV-2 is released from nerve endings. For simplicity, our model comprises a two-dimensional lattice consisting of tightly packed square cells with 8 potential cellular contacts at edges and corners: this assumption is in accordance with the arrangement of squamous epithelial cells within tissue. Many cells in the top layers of human skin are not permissive to HSV replication, so lateral cell-to-cell spread of HSV-2 is more efficient than apical to basal spread (*39*). Nevertheless, by neglecting the vertical dimension of cells, the model may underestimate total number of infected cells and viral load.

To approximate the observation that infection of adjacent cells occurs via tight-junctions,(*40*) we allow individual viruses to diffuse with random directionality across the grid immediately after their production. Viruses that reside in the same model region as susceptible cells generate infection with a concentration-dependent probability at each time step **(Materials and Methods)**.

### Unimpeded HSV-2 spread in the absence of T_RM_

In simulations without T_RM_, viral load expands rapidly during the first 12-18 hours of infection **(Fig. 3b)**, but reaches maximal values lower than those observed during the most severe shedding episodes **(Fig. 1a**,**b)**. This discrepancy is likely due to our model’s two-dimensional structure of cells. Moreover, the empirical data often include the sum of viral contributions from multiple ulcers, whereas we only model one ulcer (*19*).

In the absence of T_RM_ activity, viral load does not peak but instead grows asymptotically with deceleration after one day **(Fig. 3b)**. Both total number of infected cells **(Fig. 3c)** and ulcer diameter **(Fig. 3d)** expand in an unimpeded fashion. Realistic estimates of ulcer diameter (1-3 mm) are noted at 24-48 hours following infection with large ulcers after several days **(Fig. 3d)**.

A two-dimensional structure of uncontrolled infection emerges from the model **(Movie S1)**. The leading edge of simulated ulcers consists of pre-productive, newly infected cells, surrounding a thicker band of productively infected cells and a central core of dead and regenerating cells **(Fig. 3e)**. An outer ring of high viral load is noted after two days **(Fig. 3f)** with a far outer ring of lower viral load due to viral diffusion. The asymptotic behavior of viral load relates to spatial features of the mathematical model: target cell limitation increases with further expansion of the simulated ulcer. Specifically, as infection spreads radially, new infected cells are increasingly likely to contact already infected or dead cells (*41*).

Persistent high viral load shedding and large non-healing ulcers are well-described features of infection in persons with severely compromised T cell immunity due to HIV/AIDS or stem cell transplantation(*42*). Our basic model recapitulates untreated HSV-2 infection in these clinical populations.

### A mathematical model incorporating T_RM_ trafficking, dendricity, local proliferation, and rapid contact-mediated killing of infected cells

Most immunocompetent persons start to clear each HSV-2 shedding episode within a day after viral load peaks **(Fig. 1)**. To capture this fact, we added T_RM_ immunity to the baseline model (**Materials and Methods**, **Table 2**). For each simulated episode, we generated varying realistic densities of T_RM_ across the modeled grid based on our analysis of biopsy specimens **(Fig. 2)**, by randomly sampling rank abundance slope and T_RM_ abundance within the highest ranked region **(Fig. 2d)**. Forty percent of cultured CD8+ T cells isolated from HSV-2-infected tissue are activated in the presence of whole virus(*9*); in our simulations, this percentage of T_RM_ retains the ability to recognize HSV-2 antigen and kill adjacent infected cells in a contact-dependent manner.

**Table 2.** T_RM_ parameters

| Parameter | Unit | Value | Source |
| --- | --- | --- | --- |
| T<sub>RM</sub> starting density | - | varied | (20) |
| HSV-specific fraction<sup>1</sup> | - | 0.4 | (9) |
| T<sub>RM</sub> motility | sites day<sup>-1</sup> | 24 | (1, 22) |
| Death rate (Ag-)<sup>2</sup> | day<sup>-1</sup> | 0 | (65) |
| Proliferation (Ag+) | day<sup>-1</sup> | 3 | (65, 66) |
| Killing rate<sup>3</sup> | day<sup>-1</sup> | 48 | (25) |
| Max T cell divisions | - | 5 |  |

1 Fraction of starting T<sub>RM</sub> that are HSV-specific, remaining are bystander T<sub>RM</sub>
2 T<sub>RM</sub> death rate in the absence of antigen
3 From antigen detection to death of infected cell, per CTL pairing

Numerous recent murine studies allow parameterization of T_RM_ behavior in tissue during the immunosurveillance phase between HSV-2 shedding episodes. When cognate antigen is not present, T_RM_ patrol the cellular grid (**Fig. 4a**) (*43*). Recent evidence provides rates for T_RM_ movement and suggests that trafficking follows a random walk over time scales of hours (*1, 22*). Patrolling T_RM_ project dendritic arms to achieve a greater number of contacts with adjacent cells (**Fig. 4a**) (*1, 22*). We selected a value for T_RM_ dendricity that captures the average number of target cells that a single T cell touches.

**Fig 4.**
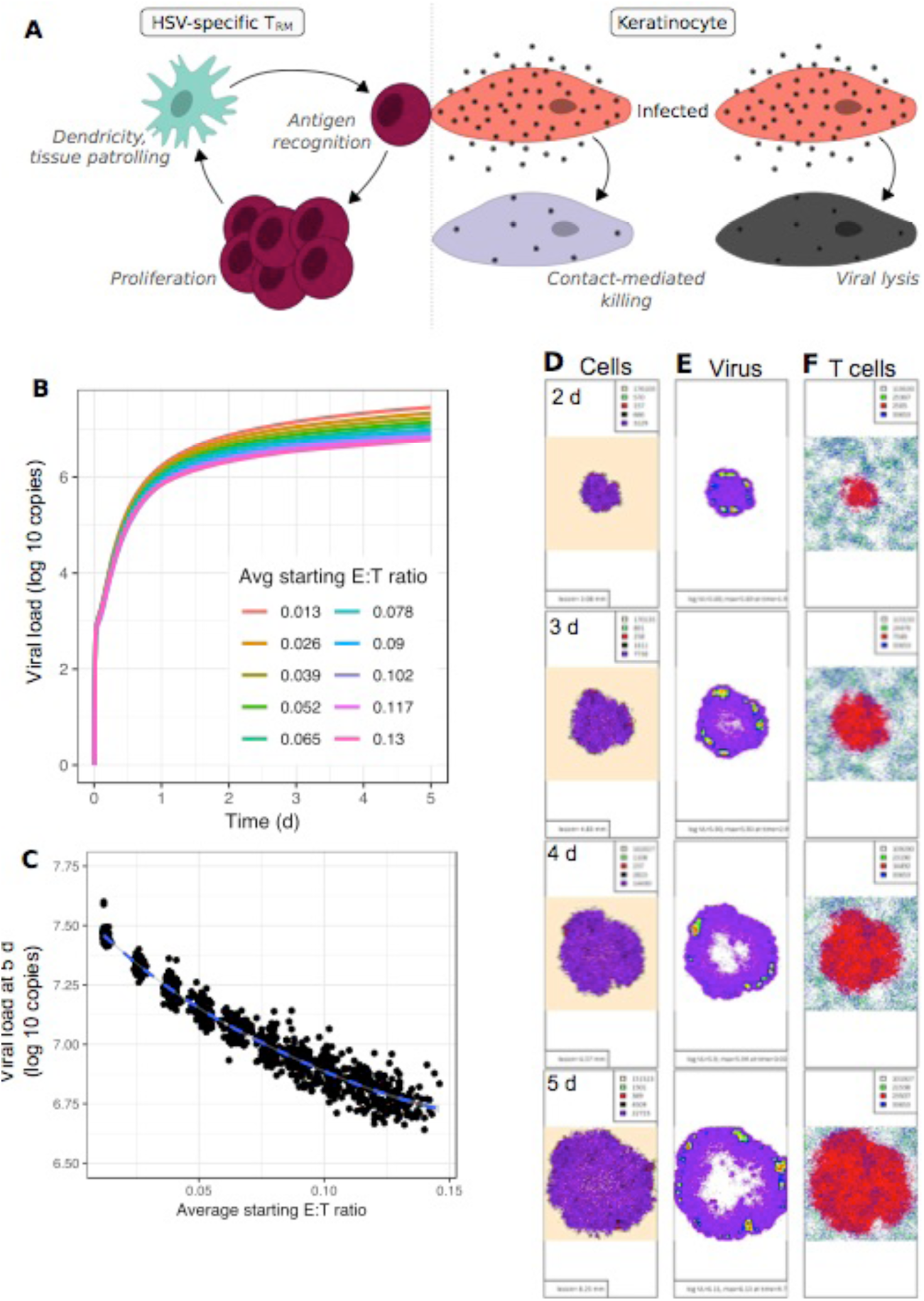
Failed elimination of infected cells at low E:T ratios in mathematical model simulations in which T_RM_ only exert contact mediated killing. **(A)** Model schematic: between HSV-2 shedding episodes, T_RM_ immunosurveillance by patrolling and expression of dendritic arms; during episodes, local T_RM_ proliferation and killing of adjacent infected cells. **(B-C)** Simulated episodes at different E:T ratios. **(B)** Viral load trajectories with asymptotic behavior and lower setpoints at higher starting E:T ratios. **(C)** Inverse correlation between E:T ratio and viral load at day 5 (dot = a simulation, n = 1000). **(D-F)** Spatial infection at high E:T ratios in 5-day simulations which correspond to **Movie S2**; one-day time steps per row. **(D)** Pre-productively infected cells (green) at the leading edge with a limited inner ring of productively infected cells (red), a core of virally lysed cells (black) and mostly T_RM_ lysed cells (purple cells). **(E)** Active HSV-2 replication within isolated intense foci of high local viral loads over regions with actively infected cells; viral loads (HSV DNA copies) per cellular region: purple=10^2^-10^2^.^25^, blue=10^2^.^25^-10^2^.^5^, green=10^2^.^5^-10^2^.^75^, yellow=10^2^.^75^-10^3^, orange=10^3^-10^3^.^25^, red=10^3^.^25^-10^3^.^5^, dark red>10^3^.^5^. **(F)** HSV-2 specific T_RM_ expansion during lesion spread (green = inactivated HSV-specific T_RM_, red = activated HSV-specific T_RM_, blue= inactivated bystander T_RM_).

HSV-2 specific T_RM_ undergo phenotypic changes upon contacting and engaging an infected cell, taking on a round morphology and transiently limited mobility. Brisk *in situ* T_RM_ proliferation ensues, increasing T_RM_ levels approximately 10-fold over a 24-hour period (**Fig. 4a**) (*23, 24*). Most local amplification of local T cell levels within 72 hours of antigen introduction is via T_RM_ proliferation rather than trafficking of CD8+ T cells from blood or draining lymph nodes (*23, 24*). In most cases, HSV-2 DNA has been eliminated from tissue at this point. Therefore, we included *in situ* T_RM_ proliferation in our model, but excluded systemic trafficking of T cells.

Upon recognizing and adhering to an infected cell, T_RM_ induce apoptosis within 30 minutes, presumably via secretion of granzyme B and perforin **(Fig. 4a)** (*25*). In our simulations, T_RM_ occupy sites on the grid of epidermal cells. If T_RM_ are at sites adjacent to (via a corner or edge), or in the same location as, an infected cell, they can kill these infected cells via TCR contact-mediated mechanisms at a fixed rate or in a synergistic fashion, such that an infected cell contacted by several T cells dies more rapidly **(Materials and Methods)** (*25*). We also assume that a single T cell can concurrently kill multiple adjacent infected cells via dendritic extensions (*44*). Cytokine paracrine effects against infected cells were not included in this version of the model.

### Unimpeded HSV-2 spread in the presence of T_RM_ that exert contact-mediated killing but no paracrine effects

When we simulated the model with the above assumptions of T_RM_ immunosurveillance and pathogen containment, there was insufficient immunologic pressure to eliminate all infected cells. Even at extremely high E:T ratios (>0.20) rarely observed in our specimens (**Fig. 2e)**, only 21% of 100 simulated episodes were contained in three days. We simulated 100 episodes at 10 realistic E:T ratios (range: 0.013-0.13). Asymptotic viral shedding was observed in all cases **(Fig. 4b).** E:T ratio was predictive of viral load setpoint in these simulations **(Fig. 4c)**, indicating that contact-mediated killing by T_RM_ exerts meaningful immunological pressure.

A spatial approximation of this infection model demonstrates that more infected cells are killed by T_RM_ than direct viral lysis **(Movie S2**, **Fig. 4d)**, and that viral loads are lower per cellular region over time **(Movie S2**, **Fig. 4e)**, relative to simulations without T_RM_ **(Movie S1**, **Fig. 3e**,**f).** Nevertheless, HSV-2 spreads effectively through micro-regions of both low and high T_RM_ density **(Movie S3**, **Fig. 4f)**. We conclude that despite the addition of multiple T_RM_ functions to our model, HSV-2 spread outpaces direct cell-to-cell killing via T_RM_ and does not recapitulate observed shedding episode features in immunocompetent humans **(Fig. 1a)**. This observation led us to include non-contact mediated functions of T_RM_ in our model.

### Secretion of granzyme B and IFN-γ early during human HSV-2 shedding episodes

To validate inclusion of cytokines in our mathematical model, we performed local sampling for cytokine quantitation every 3 hours during naturally occurring shedding episodes in persons with chronic HSV-2 infection. During all episodes, we noted an early 1-2 log surge in granzyme B **(Fig. 5a**, **upper row)** compatible with direct contact-mediated killing of infected cells by local T cells and NK cells. We detected an increase of similar magnitude in IFN-*γ* concordant with episode peaks **(Fig. 5a**, **lower row)**, suggesting that T_RM_ exert paracrine effects. Because CD8+ T cells outnumber CD4+ T cells and NK cells in HSV-2 lesions at the dermal-epidermal junction (*16*), these cells are likely to be critical producers of both detectable granzyme B and IFN-*γ* though we cannot rule out contribution from CD4+ T cells and NK cells in the dermis. Keratinocyte derived cytokines including IFN-α and IFN-β did not consistently surge during these episodes and were often absent or present at much lower levels **(Fig. S1a**, **b)**.

**Fig 5.**
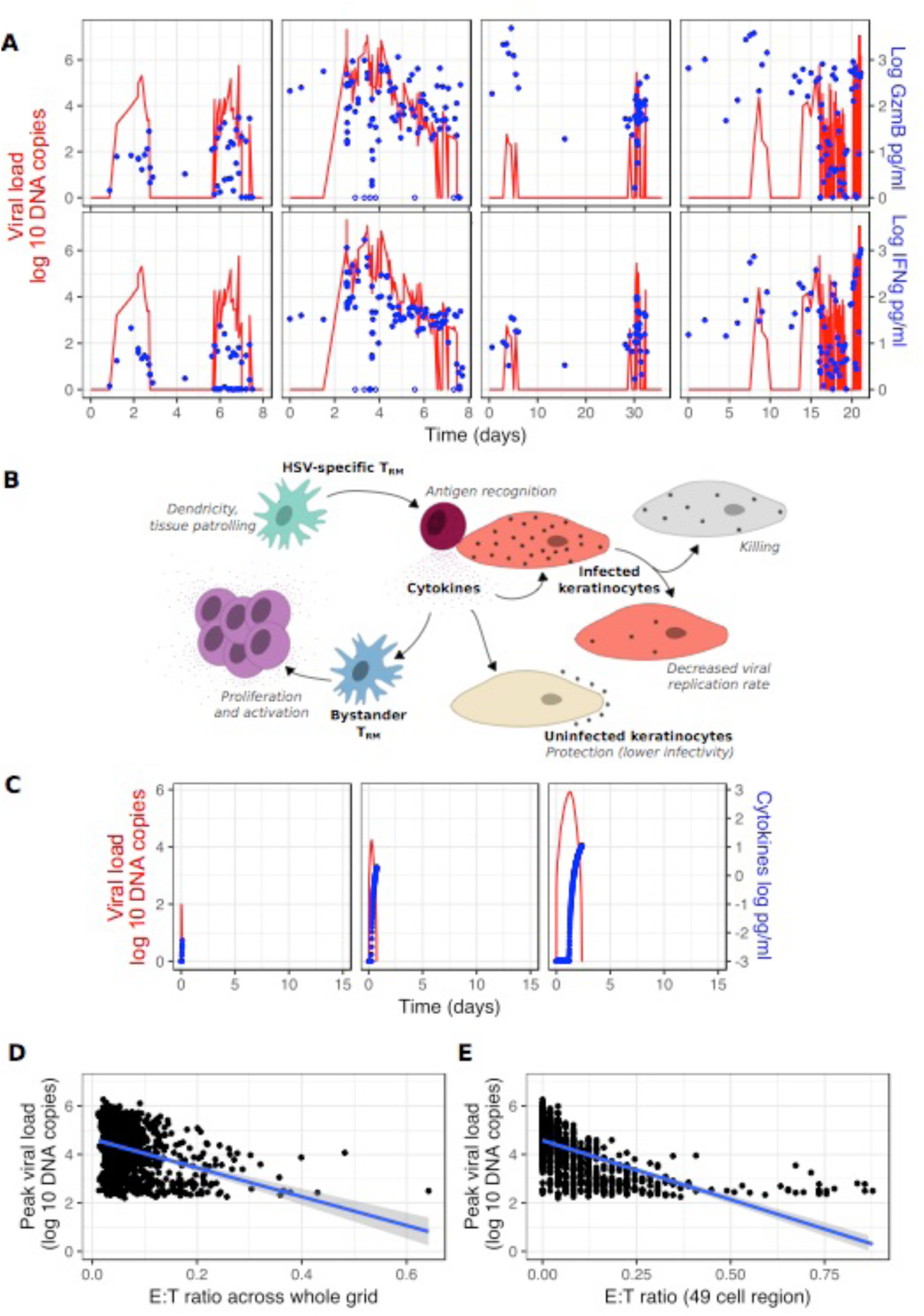
T_RM_ induced cytokine surge leading to rapid elimination of infected cells. **(A)** HSV-2 shedding episodes in four infected persons who swabbed every 8 hours pre-lesion detection and every 3 hours post-lesion detection; equivalent episodes aligned vertically; red line = HSV-2 DNA (left y-axis). **(A**, **top row);** surge in granzyme B (blue dots) during viral expansion indicating T-cell mediated killing directly via the T cell receptor. **(A**, **bottom row);** surge in IFN-γ levels (blue dots) indicating secretion and paracrine effects by local T_RM_. **(B)** Mathematical model schematic: possible cytokine effects including enhanced killing of infected cells, lowering viral replication rate in infected cells, lowering HSV-2 infectivity to uninfected cells and activating other bystander T_RM_ to proliferate and secrete cytokines; the complete model, including all cytokine functions, best fit to the data **(Fig. S2) (C)** Three simulated episodes starting at different E:T ratios; log 10 HSV-2 DNA (red) and relative cytokine concentration (blue). **(D**,**E)** 1000 simulated episodes with starting E:T ratio densities randomly selected from **Fig. 2d;** inverse correlation of the ratio of T_RM_ to epithelial cells within **(D)** the entire model grid of 15625 cells and **(E)** the 49 cells surrounding the first infected cell to peak viral load.

### Model refinements to assess possible cytokine mechanisms for containment of local HSV-2 infection

T_RM_-secreted cytokines may generate a local alarm state via numerous paracrine mechanisms after diffusing across a field of cells. We updated our model to include four potential cytokine functions: 1) enhanced elimination of infected cells, 2) reduction in viral replication rate, 3) reduction in HSV-2 infectivity and 4) induction of bystander (non HSV-specific) T_RM_ proliferation and cytokine production **(Fig. 5b)**. Inclusion of these mechanisms is experimentally justified: T_RM_-derived cytokines, particularly IFN-*γ*, induce apoptosis in HSV-2 infected cells(*27*), and lower viral replication rate. IFN-*γ* renders keratinocytes resistant to HSV-2 infection for several days in mice(*2*). Finally, cytokines may stimulate proliferation, differentiation and innate-like cytokine secretion from other local bystander CD8+ T cells in a paracrine fashion (*45-49*).

Because it is unknown which of these mechanisms are relevant during human infection, we generated 16 models with every possible combination, including a model lacking any cytokine functionality and one with all four functions. For all models, we assumed that bystander T_RM_ are incapable of recognizing and lysing infected cells via direct contact.

We imputed experimentally observed values of cytokine production by activated T_RM_ into the model **(Table 3)**. The rate of cytokine diffusion across a cellular scaffold was estimated *in vitro*(*29*), and exceeds that of viral cell-to-cell spread. Given that rates of cellular uptake and diffusion of cytokines *in vivo* are unknown, we varied these two parameters across wide ranges of values for all models tested. For models including one or more of the mechanisms described above, we varied additional parameters. For instance, we included values to characterize the cytokine concentration at which a specific cytokine action was half-maximal in a given region.

**Table 3.** Cytokine parameters

| Parameter | Unit | Value | Source |
| --- | --- | --- | --- |
| Secretion by T<sub>RM</sub> | pg cell<sup>-1</sup> day<sup>-1</sup> | 0.02 | (67) |
| Diffusion rate | sites day<sup>-1</sup> | 65 | Fitted |
| Uptake by cells | pg cell<sup>-1</sup> day<sup>-1</sup> | 6.9 | Fitted |
| Decay rate | site<sup>-1</sup> day<sup>-1</sup> | 0.01 | Fitted |
| IC50: infected cell killing | pg cell<sup>-1</sup> | $5.4 \times 10^{-6}$ | Fitted |
| IC50: viral infectivity | pg cell<sup>-1</sup> | $7.0 \times 10^{-7}$ | Fitted |
| IC50: virus production | pg cell<sup>-1</sup> | $1.0 \times 10^{-6}$ | Fitted |
| IC50: T<sub>RM</sub> activation | pg cell<sup>-1</sup> | $1.6 \times 10^{-7}$ | Fitted |

We explored 1000 possible randomly selected parameter sets for the 16 models. For each unique parameter set, we simulated 100 episodes, each with different initial conditions of T_RM_ density drawn randomly from exponential distributions **(Fig. 2d)**. In total, we simulated 1,600,000 HSV-2 ulcers.

### Recapitulation of observed shedding episode kinetics with incorporation of HSV-2 specific T_RM_ cytokine secretion into the mathematical model

To identify which cytokine functions and parameters best recapitulate the observed data, we performed comparative model fitting to shedding episode data using Akaike Information Criteria (AIC), to reward for fit while penalizing for unnecessary complexity. We fit to shedding data from the 83 episodes described in **Fig. 1**. Episodes were arrayed in rank abundance curves according to peak viral load, duration and time to peak, such that the models were tasked with reproducing the entire distribution of shedding episode severity **(Fig. S2).** To mimic the clinical protocol, we assumed sampling every 6 hours and random episode initiation 0 to 6 hours after each simulated sample.

Optimal data fitting occurred with inclusion of all four cytokine mechanisms in **Fig. 5b (Fig. S2a)**. We added a fifth possible mechanism to the model, cytokine induction of a chemical gradient to allow CD8+ T cells to home to sites of infection: this did not enhance data fitting and was not subsequently included. Similarly, adjusting modes of T_RM_ trafficking from random walk to Levy walk did not alter model fit and was not subsequently included. These results suggest that elimination of HSV-2 infected cells by cytokines involves multiple complementary mechanisms.

The optimized model slightly overestimated distributions of episode peak viral load and durations at moderate ranks **(Fig. S2b,c)**, underestimated peak viral load of the most severe episodes **(Fig. S2b)**, but closely recapitulated distribution of time to episode peak **(Fig. S2d)**. The model accepted a wide range of plausible values for rates of cytokine uptake into cells and viral diffusion. A narrower range was identified for cytokine diffusion rate, indicating the importance of this value for model fitting **(Fig. S2e)**. Threshold upper limits were identified for necessary intracellular concentration of cytokines to limit viral replication and induce death in infected cells, prevent infection in uninfected cells, and activate surrounding T_RM_. These values were low, suggesting that cells are extremely sensitive to these proteins (**Table 3**).

### Rapid containment of infection under low T_RM_ condition due to rapid cytokine diffusion derived from HSV-2 specific T_RM_

Viral load trajectories in simulated episodes were notable for early peaks, rapid expansion rate and short duration **(Fig. 5c)**; timing of cytokine surge **(Fig. 5a, c)** was comparable to human shedding episodes. As with murine studies (*24*), E:T ratio in the infected area was fairly predictive of peak viral load in simulations: this was true when a field of 15625 or 49 target cells was considered **(Fig. 5d, e)**. A T_RM_ to epithelial cell ratio of roughly 0.2 was required for rapid elimination of HSV-2 infected cells prior to development of viral loads sufficient for lesion development and sexual transmission potential (>10^4^ HSV DNA copies) **(Fig. 5d, e)**; this exceeded ratios observed in most human biopsy specimens **(Fig. 2e)**. Early recognition of infection by T_RM_ often resulted in elimination of virus in 6-24 hours with fewer than 20 total infected cells **(Movie S4).**

Spatial visualization demonstrates that in more severe episodes, later initial recognition of an infected cell by an HSV-2 specific T_RM_ (due to a lower T_RM_ density) is followed by rapid diffusion of cytokines and immediate impact of cytokine antiviral effects **(Fig. 6a**, **Movies S5 & S6)**. Accordingly, in simulations the time between infection of the first cell and recognition of an infected cell by an HSV-2 specific T_RM_ was predictive of peak viral load **(Fig. 6b)**, number of infected cells **(Fig. 6c)** and episode duration **(Fig. 6d)**. As episodes increased in severity, the proportion of infected cells killed by cytokine effects rather than direct lysis by T_RM_ was generally higher **(Fig. 6e)**.

**Fig 6.**
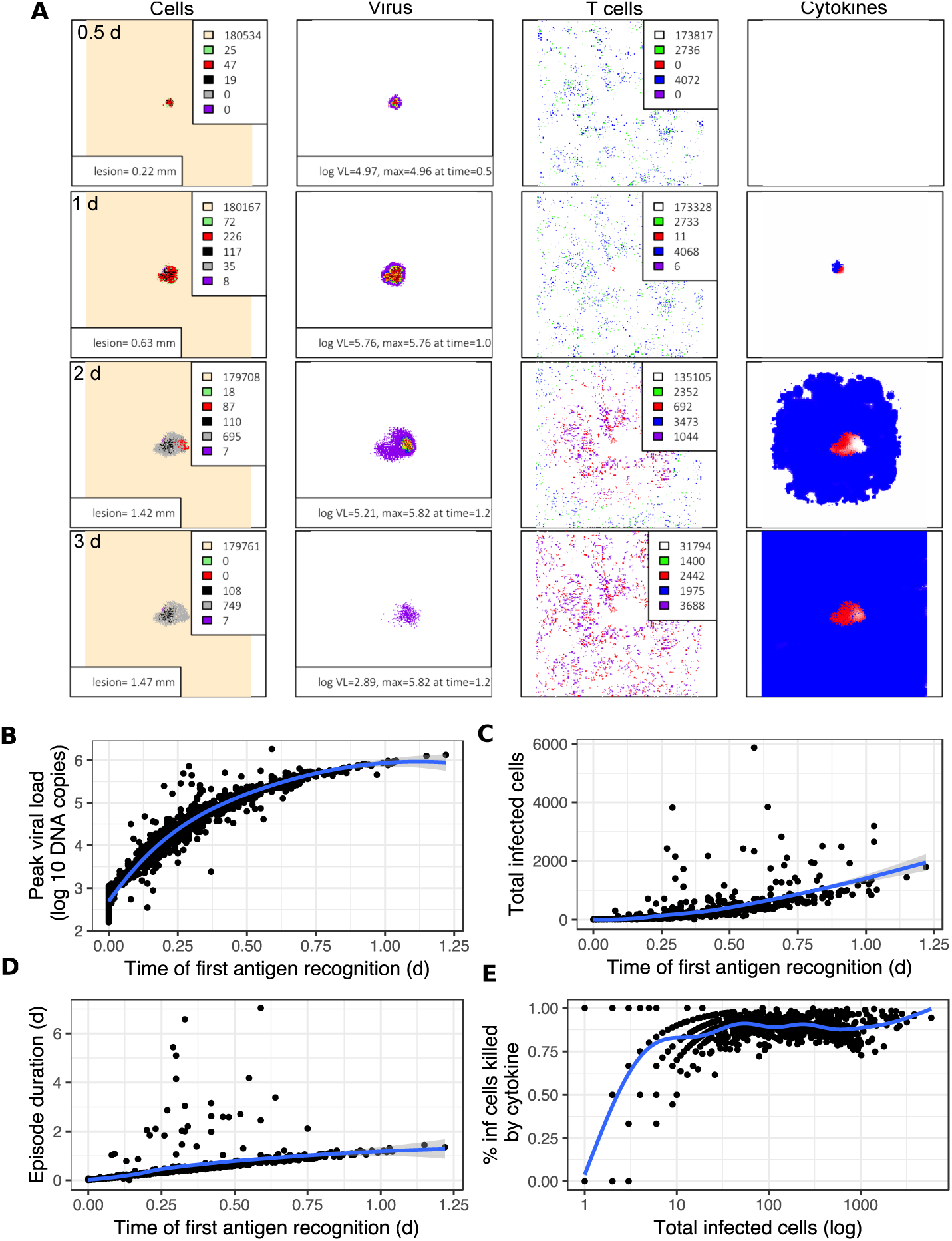
Cytokine spread leading to rapid elimination of infected HSV-2 cells. **(A)** A simulated episode (corresponding to **Movies S5 & S6)** with low T_RM_ density leading to control of infection. Panels left to right: (1) epidermal cell dynamics, peach = uninfected, green = pre-productive infection, red = productive infection, black = virally lysed cell, purple = T_RM_ lysed cell, grey = cytokine lysed cell; (2) viral loads (HSV DNA copies) per cell: purple=10^2^-10^2^.^25^, blue=10^2^.^25^-10^2^.^5^, green=10^2^.^5^-10^2^.^75^, yellow=10^2^.^75^-10^3^, orange=10^3^-10^3^.^25^, red=10^3^.^25^-10^3^.^5^, dark red>10^3^.^5^; (3) T_RM_, green = inactivated HSV-specific T_RM_, red = activated HSV-specific T_RM_, blue = inactivated bystander T_RM_, purple = activated HSV-specific T_RM_; (4) Concentration of cytokine by color darkness; red over infected or dead cells, indicating reduced viral replication and infected cell lifespan; blue cover uninfected cells, indicating reduced viral infectivity. **(B-E)** 1000 simulated episodes with initial E:T ratio densities randomly selected from **Fig. 2d.** Correlations of time to first T_RM_ recognition of an infected cell with **(B)** log10 peak viral load, **(C)** total infected cells, and **(D)** duration. **(E)** Proportion of infected cells killed by cytokine effects as a function of number of total infected cells per episode. Blue lines = LOESS-smoothed lines with 95% CI in grey.

### Polyfunctional and individually redundant antiviral cytokine effects within the HSV-2 infection micro-environment

To further delineate mechanisms of cytokine protection, we performed *in silico* knock out experiments by removing single and multiple cytokine mechanisms over 100 sequential shedding episodes. These simulations demonstrated that elimination of single cytokine mechanisms generally had a limited effect on episode severity; yet, when multiple cytokine mechanisms were not included in the model, a higher proportion of episodes had longer time to peak viral load **(Fig. S3a)**, higher total number of infected cells **(Fig. S3b)**, higher peak viral loads **(Fig. S3c)** and longer duration **(Fig. S3d)**. Among cytokine mechanisms, activation of bystander T_RM_ (particularly induction of cytokine secretion) and limitation of infected cell lifespan were most critical in limiting episode duration. Simulations in which cytokines did not impact infected cell lifespan **(Movie S7)** or HSV infectivity **(Movie S8)** led to slightly protracted episodes in which ulcers took on an uncharacteristic serpiginous appearance.

### More rapid HSV-2 elimination due to T_RM_ trafficking, dendricity, proliferation, and cooperative killing

We performed additional *in silico* knock out experiments to identify the relative importance of non-cytokine mediated T_RM_ mechanisms of immune control. Elimination of single immunosurveillance mechanisms generally had a limited effect on episode severity; when both T_RM_ dendricity and mobility were eliminated from the model, a higher proportion of episodes had longer time to peak viral load **(Fig. S4a)**, higher total number of infected cells **(Fig. S4b)**, higher peak viral loads **(Fig. S4c)** and longer duration **(Fig. S4d)**. Elimination of contact-mediated killing had no overall effect on episode severity, highlighting the potency of antiviral cytokine spread.

## Discussion

T_RM_ play a critical role in mucosal protection against HSV-2. During chronic infection, genital tissue contains HSV-2 specific and bystander T_RM_ months after local elimination of infected cells (*9*). The release of HSV-2 from latently infected ganglia via sensory neurons into genital tissues is temporally and spatially stochastic, resulting in seeding of low and high density T_RM_ regions (*19, 20, 50*). Within micro-regions, the spatial heterogeneity of T_RM_ results in highly variable amounts of viral replication and subsequent T_RM_-mediated expansion (*21*). At the whole tissue level, the result is a shedding pattern characterized by some brief episodes (approaching sterilizing immunity) in which infection is limited to a few cells, and other episodes that persist for days and are associated with lesions, infection of thousands of cells and high transmission risk (*51, 52*).

While HSV-2 achieves sufficient replication to induce a disease state in many infected people, the immune system wins each local battle with remarkable efficiency. Viral loads invariably peak within 24 hours after viral DNA detection; the abrupt transition from viral expansion to contraction is associated with a surge in granzyme B and IFN-*γ*. Using a mathematical model, we demonstrate that HSV-2 specific T_RM_ induce rapid apoptosis of directly adjacent infected cells. If this occurs when infection is limited to one or a few cells, then the episode will terminate without need for an amplified local response. If the number of infected cells is greater, then T_RM_ contact-mediated killing is insufficient for pathogen control, despite efficient T_RM_ trafficking, expression of dendritic arms to canvas a high number of local cells, *in situ* proliferation, and rapid cooperative killing of infected cells.

The essential second step for truncating local shedding is cytokine secretion, which leads to a profound antiviral state in surrounding cells. Because they are smaller, secreted cytokines diffuse more rapidly than viruses, and decelerate the pace of HSV-2 spread (*29*). Based on best fit to observed data, our model suggests that cytokines exert distinct effects including rapid killing of infected cells, lowering HSV-2 replication, lowering HSV-2 infectivity to uninfected cells, and activating bystander T_RM_. Experimental evidence suggests that IFN-γ mediates protection by reducing HSV-2 infectivity to susceptible cells or increasing apoptosis of infected cells (*26, 27*). Inclusion of the latter mechanism in our model was crucial for recapitulating early viral load peak. The rapid lethality of diffusing cytokines explains the abrupt formation of clinical vesicles and erosions concurrent with viral clearance.

The generalized cytokine alarm state mechanism explains how HSV-2 elimination occurs rapidly within mucosal micro-regions defined by low HSV-specific T_RM_ density (*2*). In our model, recognition of infected cells by a single pathogen-specific T_RM_ is necessary but insufficient for initiation of tissue-wide protection. The timing of this sentinel event during the early hours of infection determines the extent of viral replication.

As was demonstrated in a mouse model, the total density of HSV-2 specific T_RM_ is likely to be a key surrogate of immunity (*24*). At a scale of a 3 mm × 3 mm simulated micro-region, or a smaller 50-cell region, T_RM_ density maintains an inverse dose response with both peak viral load and shedding duration. Therefore, to achieve full protection, a therapeutic vaccine must achieve high HSV-2 specific T_RM_ levels within all micro-regions across the genital region. During natural infection, the number of HSV-2 specific T_RM_ is only sufficient to provide rapid protection in approximately 20% of tissue (*20*).

The model projects that bystander T_RM_, which cannot recognize infected cells, but secrete cytokines upon activation, play an important role in late elimination of infected cells during severe episodes. However, a high density of T_RM_ that lack antigenic specificity is not predicted to be a surrogate of local immune protection. A high E:T ratio of HSV-2 specific cells is the key requirement for near sterilizing immunity.

Our *in silico* knock out analysis allows a detailed assessment of the relative contribution of T_RM_ phenotypic characteristics to the pace of HSV-2 elimination. Single functions can typically be eliminated without dramatically impairing control of infection. For instance, elimination of trafficking or T_RM_ dendricity has only modest effects on viral load during simulations; however, elimination of both functions leads to a substantial increase in episode duration and peak viral load. Cytokine effects also have substantial redundancy.

Our approach has important limitations pertaining to oversimplification of the totality of the immune response. We are unable to discriminate the effects of CD4+ and CD8+ T_RM_,(*28*) and do not consider infiltrating natural killer cells (NK cells). CD8+ T cells co-localize with infected cells at the dermal-epidermal junction to a greater extent than CD4+ T cells which provide help at a distance from the dermis (*16*). NK cells lack antigen specificity and are present in lower abundance in healing genital biopsy specimens (*8, 16*). Nevertheless, both cell populations are vital for control of human herpes viruses, exhibit memory, and secrete IFN-*γ*, perforin and granzyme B (*10, 53*). Therefore, NK cells and CD4+ T_RM_ may serve as bystander cells and contribute to the general alarm state (*8*). It will be important to distinguish the role of various infiltrating immune cell subsets in future models as relevant data accrues.

We do not include epithelial cell-induced innate immunity, antigen presenting cells (*54*), or antibody effect (*55*). While these mechanisms play a role in pathogen elimination, to our knowledge their effects may not be as variably dispersed over time and tissue space as T_RM_. Therefore, our model may capture humoral and local innate immunity effects which are embedded within static model parameters for viral replication, infectivity and spread; if these effects vary in a non-linear fashion relative to local viral load, then our model misses these dynamics. During some contained episodes, study participants did not mount a robust IFN-α and IFN-β response, indicating that epithelial-derived innate responses are sometimes suppressed. Yet, during severe episodes, it is possible that infected epithelial cells and antigen presenting cells sufficiently activate bystander T_RM_ even in the absence of HSV-specific T_RM_.

We denote cytokines as one variable in our model, which is overly reductionist. The multiple antiviral effects of IFN-*γ* are crucial, but a complex network of cytokines is likely to influence HSV-2 containment, with varying effects according to cellular source and cellular target. While a powerful cytokine impact on infected cells is a robust feature of our model, a more comprehensive assessment will only be possible as new data allows detailed parameterization of individual cytokine effects.

Our model projections deviate slightly from the observed data. The best fitting model does not achieve the highest peak viral loads, possibly because the two-dimensional model includes fewer target cells for HSV-2 replication than human skin. Our model also only simulates a single ulcer while swabbing protocols capture HSV-2 from multiple contemporaneous ulcers when viral loads are highest (*19*). The optimal model slightly overestimated the distribution of episode durations and peak viral loads at moderate ranks, perhaps because other infiltrating immune cell populations not included in the model contribute to control of infection.

Despite these limitations, modeling provides a unique platform to link observations from murine and human experiments, which is important because both of these approaches have inherent gaps. In humans, T_RM_ responses evolve quickly within microscopic tissue environments and have not been captured during critical early stages of viral replication and spread. Successful containment of HSV-2 in a few cells, is asymptomatic, and even more difficult to observe empirically. Yet, quantitative components of human data, including heterogeneity in shedding episode kinetics and spatial density and topology of T_RM_ serve as critical benchmarks for model validation.

Murine models utilizing intravital imaging allow observation and quantification of T_RM_ behaviors. However, these experiments are rarely centered around realistic kinetics of pathogen control. Moreover, murine infection conditions may not be representative of human HSV-2 reactivation. It is plausible that T_RM_ trafficking, dendricity, proliferation, synaptic killing and antiviral cytokine release occur in humans and mice. Yet, *in vivo* rates governing these behaviors may differ. Our analysis shows that knock out of many of these mechanisms results in perturbations in viral shedding outcomes.

HSV-2 infected individuals experience shedding episodes that are frequent and highly variable due to the spatial distribution of T_RM_. We propose that containment of infection within low T_RM_ density areas is contingent upon rapid diffusion of antiviral cytokines from activated T_RM_. Effective vaccine strategies must increase the total number of HSV-2 specific T_RM_ across the entire genital tract and leverage the broad protection afforded by antiviral cytokines to induce a spatially homogeneous layer of immunological readiness.

## Materials and Methods

### Study Design

We re-analyzed data from a previous study where HSV-2 seropositive, HIV-negative individuals were asked to self-collect genital swab specimens 4 times a day for 60 days. Viral load was measured using qPCR (*30*). For each participant, we divided the time series of viral loads into individual shedding episodes. We defined a single episode as a series of HSV-positive swabs with two negative swabs before and after. We discarded episodes where swabs were more than 12 h apart (i.e. 2 or more consecutive missing samples) and classified episodes as short (0 - 1 d], medium (1-2 d] and long (>2 d). For each episode, we quantified peak viral load, viral load at the first positive swab, time to first episode peak and episode duration. Episode analysis and visualization were performed in R (*56*). We also analyzed viral swab data from 3 participants for both viral load and cytokine levels (described below).

### Image analysis of human biopsy specimens

We re-analyzed RGB images from previously described human HSV-2 genital biopsy specimens showing cell nuclei (blue), CD4+ T cells (green) and CD8+ T cells (red) (*20*). Each of 10 participants was biopsied at 2 and 8 weeks after healing of the lesions. We included both biopsies in the analysis for each participant. We focused our analysis on CD8+ T cells in close proximity to the dermal-epidermal junction (DEJ), cropping all areas below the DEJ that were more than half the thickness of the epidermis below the dermal-epidermal junction. These cut areas typically had high neuronal density and low nuclear density. We enumerated CD8+ T cells and total nuclei representative of T cells and epithelial cells using the program Cell Profiler (*57*). Using a custom R script, we used the output from Cell Profiler for each image to analyze clustering of CD8+ T cells and to calculate E:T ratios within micro-regions from the biopsy. We started by dividing the image into micro-regions containing approximately 100 cells by performing k-means clustering on the positions of cell nuclei. We then assigned each CD8+ T cell to a micro-region by determining which cluster centroid was closest to each CD8+ T cell **(Fig. 2b**,**c)**. Finally we computed E:T ratios for each cluster and for the biopsy overall. Code and data are available at http://github.com/proychou/SpatialHSV.

### Cytokine data from swabs

Four study participants with recurrent HSV lesions were enrolled in an additional protocol in which swabs were obtained every 8 hours prior to lesion development. At the time of prodrome or ulcer formation, participants switched to every 3-hour sampling for 5 days. Two swabs were obtained at each timepoint. One was processed for HSV PCR (*36*), and the other was sent for MSD assay for cytokine quantitation (*58*).

### Individual-based model of viral spread and immunological containment of infection by T_RM_

We developed a stochastic, agent-based, spatially explicit model of HSV infection that incorporates key features of host and virus life cycles and immune control by CD8+ T cells, based on a previously developed model of host-pathogen dynamics (*59*). The model is run on a 2D grid of sites; each site on the L × L grid is either vacant (no cell, no virus) or occupied by a single epithelial cell and one or more virus particles. In addition, each site may be patrolled by a CD8+ resident memory T cell (T_RM_) that is either HSV-specific or a bystander in the context of HSV infection. The size of the grid (L) ranged between 125×125 and 425×425 depending on the simulation.

At each site, we track the state of the epithelial cell, the state of any T_RM_ patrolling the site, the number of infectious viruses present, and the concentration of cytokines at the site. At the start of the simulation, all sites are occupied by susceptible cells and a lawn of tissue-resident T cells at a pre-defined spatial density (defined as the probability that the focal site at the center of the grid contains T_RM_). State transitions occur stochastically at probabilities that are determined by the rates of various events that can occur at the site described below (*59*). These rates determine the wait time, a random variable whose mean is the reciprocal of the corresponding rate. Virus and cytokine concentrations decay exponentially at rates specified in **Tables 1-3**. To avoid doubly propagating particles or adding dependencies on the order of cell evaluation, the change in values occurs across the grid at the end of each time step. Diffusion is described by a 2D-Gaussian function applied over a 3×3 neighborhood as described in (*59*).

For all simulations involving model fitting, we populate the grid with a spatial distribution of T_RM_ drawn from the histologic slides from 10 participants in the human biopsy study described above (**Fig. 2a**). The E:T ratios from the image analysis above are used to generate rank order distributions of ratios for each biopsy and fit to an exponential curve of the form E:T ratio = p*e^c*rank^ **(Fig. 2d)**. The resulting distributions of the coefficients p and c are used when seeding clusters of 100 cells in the simulated cell grid prior to each simulated episode.

Each susceptible cell at site (i, j) becomes infected at rate βV_ij_, where β is viral infectivity, and V_ij_ is the number of viruses present at the site. Newly infected cells enter an eclipse phase during which they do not produce virus and then transition at rate ε to a virus-producing state, producing μ free virus per unit time. Productively infected cells die at rate δ_I_, while susceptible cells and pre-productively infected cells die at rate δ_S_. Vacant sites and sites occupied by dead cells are repopulated by susceptible cells at rate α. Each simulation starts with a single productively infected cell placed at the center of the grid and populations are recorded at 10-minute increments of simulated time for 5 days. Cell and virus parameters are listed in **Table 1**.

To this basic version of viral spread, we added immune mechanisms with features that can be switched on or off depending on the hypothesis being tested. When T_RM_ are present, they patrol the grid at a specified rate. Movement can occur by either a “random walk”, i.e. taking one step in a random direction within a 3 × 3 Moore neighborhood at a specified rate; “Levy walk”, a modified random walk where step size is drawn from a Levy distribution; or “directed walk” where the T cell moves preferentially along a cytokine gradient (if present) and towards an infected cell when present within a 5 × 5 Moore neighborhood. For most simulations, unless specified, we assumed a random walk. T cell parameters are listed in **Table 2**.

When an HSV-specific T_RM_ encounters an infected cell, movement stops and the T cell does one or more of the following: proliferates, kills the infected cell, and/or secretes cytokines. Rates of each of these processes were determined from the literature or inferred by fitting to data. The dendritic shape of T_RM_ is modelled by allowing HSV-specific T_RM_ to recognize and kill an infected cell within a 3 × 3 Moore neighborhood of its current position. *In situ* T cell proliferation is modeled by placing a daughter T cell at a randomly chosen vacant site within a 3 × 3 Moore neighborhood of the focal cell.

When activated, a T cell secretes cytokines at a defined rate and cytokines diffuse rapidly across the grid. When present at a site containing a keratinocyte, cytokines have multiple concentration-dependent effects including reducing infectivity of virus to susceptible cells, reducing viral replication rate in infected cells, and decreasing lifespan of infected cells. In addition, cytokines can activate bystander T cells that are not specific for HSV, allowing them to amplify the cytokine-mediated signal. Cytokine-related parameters are listed in **Table 3**. Cytokine effects (decreased viral replication, increased cell death rate, decreased viral infectivity and increased activation of surrounding T_RM_) are modeled with the following equation:

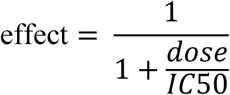

in which dose is the cytokine level (in pg/ml) within the single-cell region and IC50 is the level of cytokine level required to achieve half-maximal effect. The product is then multiplied by the parameter of relevance.

### Data fitting

We initially fit the model without T_RM_ to achieve realistically high viral loads after 24 hours of shedding. Using a search algorithm and predefined ranges from the literature **(Table 1)**, we tuned parameters of HSV-2 infectivity rate and diffusion rate, while fixing values for viral replication rate and clearance rate at experimentally observed values. We also tuned the model to diameter of the infected cell region from the empirical data. Simulated ulcer diameter was calculated from the number of dead cells (*18, 60*).

In order to evaluate models with and without various cytokine effects we used a fixed set of viral and T_RM_ parameters and varied only the parameters associated with each of the four cytokine effects. Parameter sets that did not yield at least one large episode were discarded. A large episode was defined as generating the equivalent of a 1.5 mm diameter ulcer. We fit each model to the rank order distributions of the collected cohort data described above (**Fig. 1**) for episode peak viral load, time to peak and duration. The root sum of squares (RSS) was calculated by comparing the ranked values for the cohort data with ranked values for the simulated episodes. The squared error at each point was divided by the variance of the cohort measure in order to standardize the results across the different variables. The overall AIC score was calculated using the RSS score for each category together with a complexity variable *k* that differed for each model based on the number of free parameters it introduced and a shared *n* value that represented the number of data fitting points across the three rankings. The *n* value was 83 unless the given parameter set yielded less than that number of episodes in 100 attempts. Each attempt started with one infected cell at the center of the grid. The equation used is shown below.

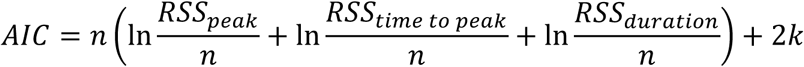

The baseline model had no cytokine effects and was assigned a *k* value of zero. The mean of the 10 highest AIC scores from parameter combinations for each of the 16 models was used to determine the best fit model (**Fig. S2a)**.

To further improve the fitting of the chosen model, we performed an additional round of simulations in which we varied viral diffusion, cytokine diffusion and cytokine uptake rates on a log10 scale across 500 parameter sets. All other parameters were held at the values for the best model. The final best parameter set from the initial fitting and fine-tuning exercise is shown in **Table 3**.

### In silico knock-out analyses

We ran simulations to evaluate the effect of removing single functions from the model and then measured results against the following metrics of episode severity: peak viral load, time to peak viral load, duration and total number of infected cells. Simulated sampling was every 6 hours to align with the cohort data in these model projections. Starting with the best parameter set for the full cytokine effects model, we ran additional simulations of 100 episodes each in which we removed one or more cytokine effects or T_RM_ features. The cytokine effects removed were reduction in viral infectivity, viral production, acceleration of infected cell death and T_RM_ activation. The latter was split into cytokine-modulated production of cytokine from bystander T_RM_ and cytokine-modulated activation of both HSV-specific and bystander T_RM_. The T_RM_ effects included reducing dendricity, halting mobility and eliminating contact-mediated killing by HSV-specific T_RM_. The resulting episodes were compared with those of the full model in several categories including average peak viral load, the total number of cells infected, episode duration and the percentage of episodes that were not controlled.

### Code

All code and data are available at http://github.com/proychou/SpatialHSV. The model is available in both R and C++ versions.

## Supplementary Materials

Fig. S1. Epithelial-derived antiviral cytokines do not predictably surge locally during HSV-2 shedding episodes.

Fig. S2. A model with all possible antiviral cytokine features optimizes fit to the data.

Fig. S3. Antiviral cytokines effects are polyfunctional and redundant within the HSV-2 infection micro-environment.

Fig. S4. Trafficking, dendricity, and contact-mediated killing, as well as activity of TRM accelerate but are not required for elimination of HSV-2 infected cells during severe episodes.

Movie S1. Lack of containment of HSV-2 spread in the absence of TRM.Movie S2. Lack of containment of HSV-2 spread in the presence of TRM which only kill via direct lysis of infected cells. Movie S3. Lack of containment of HSV-2 spread in the presence of TRM which only kill via direct lysis of infected cells.

Movie S4. Rapid containment of HSV-2 spread in the presence of TRM antiviral cytokine secretion and high TRM density.

Movie S5. Containment of HSV-2 spread in the presence of TRM antiviral cytokine secretion and low TRM density.

Movie S6. Containment of HSV-2 spread in the presence of TRM antiviral cytokine secretion and low TRM density.

Movie S7. Slower containment of HSV-2 spread in the presence of TRM antiviral cytokine secretion which only prevents infectivity.

Movie S8. Slightly protracted containment of HSV-2 spread in the presence of TRM antiviral cytokine secretion which only limit lifespan of infected cells.

## Acknowledgements

We express gratitude to our dedicated study participants, clinical research staff (Nui Pholsena, Dana Varon and Jessica Moreno), molecular laboratory scientists (Meei-Li Huang) and to helpful scientific input from Daniel Reeves, Florencia Tettamanti Boshier, and Fabian Cardozo Ojeda.

## Author contributions

P.R. conceived the study, performed mathematical modeling, performed statistical analyses, and wrote the manuscript. D.A.S. performed spatial analyses of biopsy specimens, mathematical modeling and statistical analyses. E.D. assisted with literature review and model parameter selection. L.C. conceived the biopsy study. J.Z. performed staining of biopsy specimens. V.D. and L.R.S performed cytokine analysis. J.M.L. and M.P. conceived the study and edited the manuscript. J.T.S. conceived the study, performed mathematical modeling, performed statistical analyses, and wrote the manuscript.

## Data and materials availability

All code and data are available at http://github.com/proychou/SpatialHSV.

## Supplementary Materials

**Fig S1.**
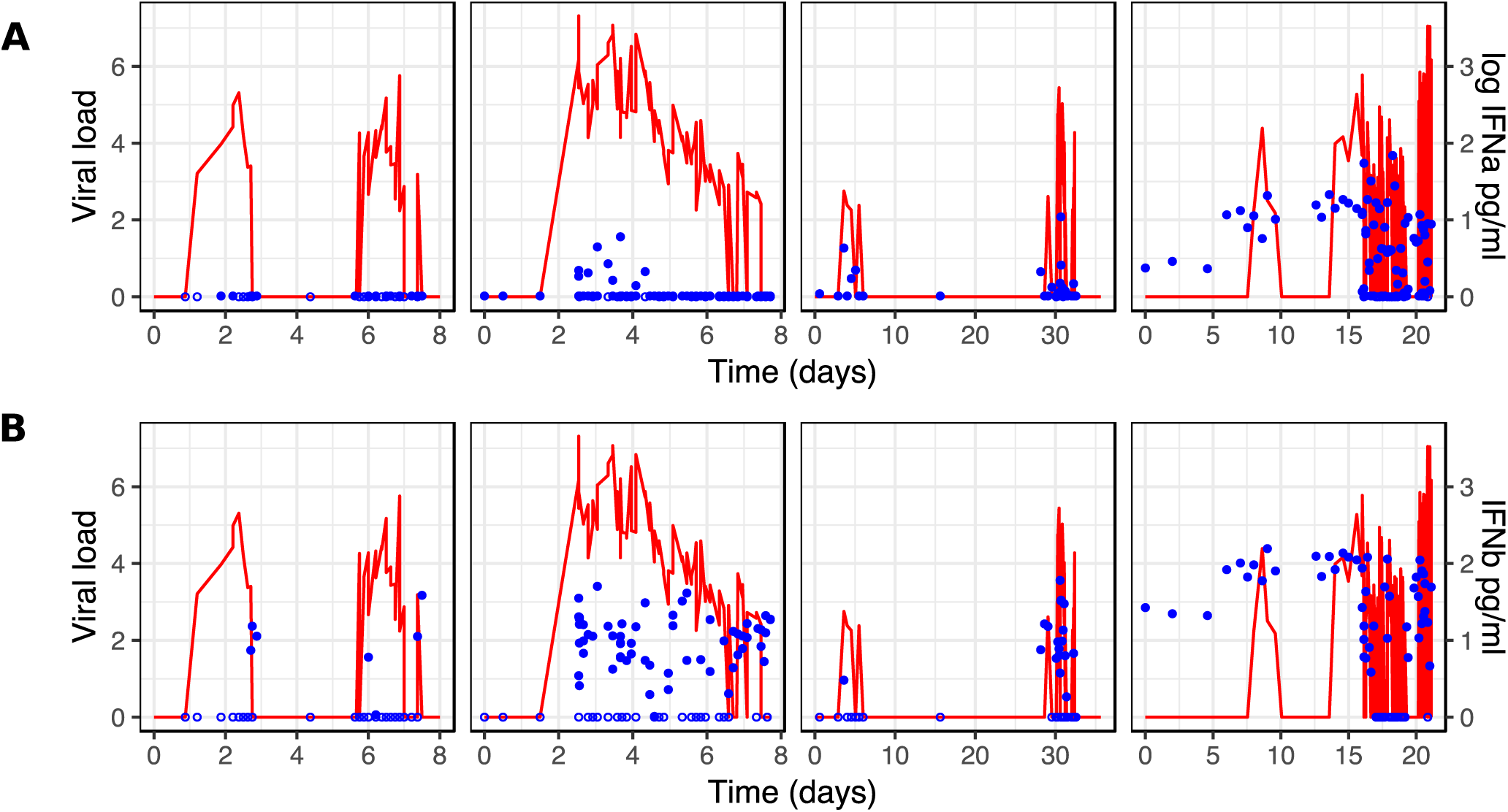
No predictable surge of epithelial-derived antiviral cytokines during HSV-2 shedding episodes. Detailed cytokine kinetics from episodes in four infected persons who performed swabs every 8 hours pre-lesion detection and every 3 hours post-lesion detection; equivalent episodes aligned vertically from **Figure 5a**; red line = HSV-2 DNA (left y-axis). **(A)** IFN-α (blue dots) during viral expansion. **(B)** IFN-β (blue dots) during viral expansion.

**Fig S2.**
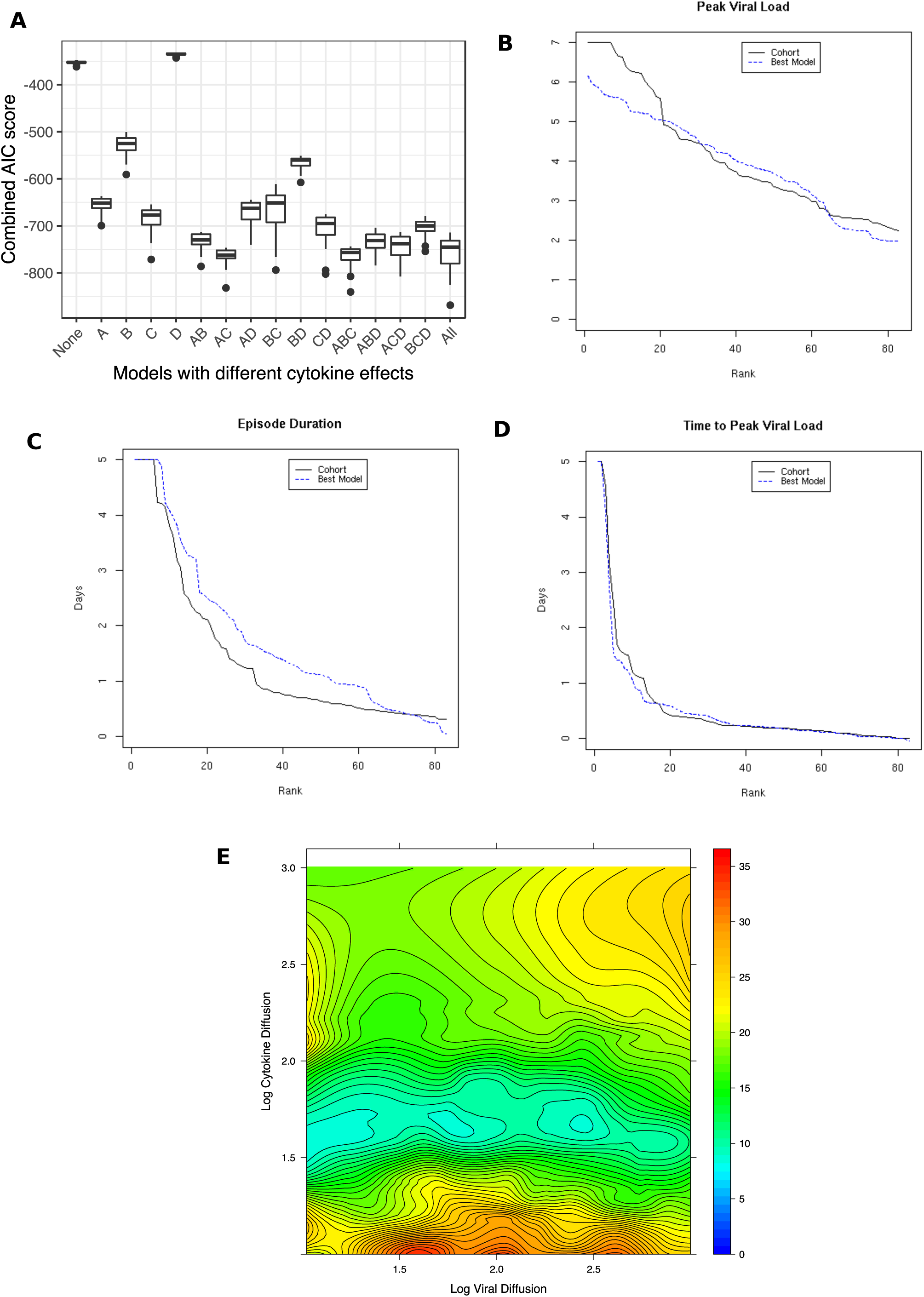
A model with all possible antiviral cytokine features with optimal fit to the data. **(A)** Optimized AIC scores for models including and excluding various possible cytokine functions including: A, increasing infected cell death rate; B, decreasing viral infectivity; C, lowering viral replication rate in infected cells; and D, activating other bystander T_RM_; low AIC scores represent higher likelihood models; lowest AIC scores occur with the most inclusive model; boxplots = interquartile ranges (IQR) and whiskers = ranges or 1.5x the IQR for 10 simulations of each model with 100 episodes per simulation. **(B-D)** Rank abundance curves of 83 episodes sorted from highest to lowest according to episode **(B)** peak viral loads, **(C)** duration and **(D)** time to peak viral load; black lines = episode data and blue lines = optimized values from the mathematical model. **(E)** Heat maps for two model parameter values (rate of viral diffusion and rate of cytokine diffusion) from the completely inclusive best-fitting model with range of values required for best fit to the data for cytokine diffusion rate and viral diffusion rate; color values = residual sum of squares between model output and the data in **B-D**: blue represents best model fit.

**Fig S3.**
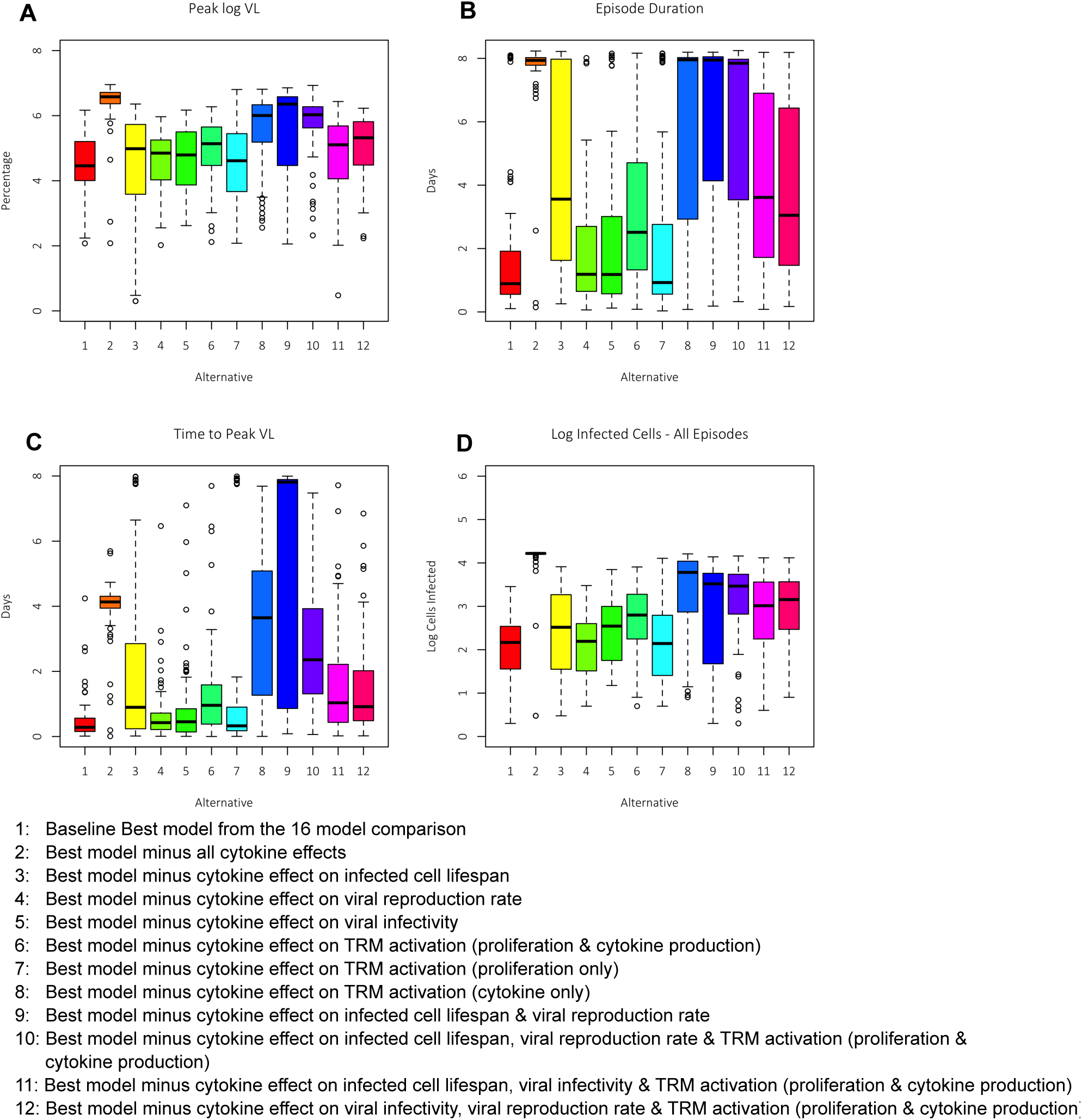
Polyfunctional and individually redundant antiviral cytokine effects within the HSV-2 infection micro-environment. Episodes simulated with starting E:T ratio densities randomly selected from **Fig. 2d** under conditions in which cytokine effects were limited to one or more of the following mechanisms: enhancing infected cell death rate, decreasing viral replication rate in infected cells, decreasing infectivity to uninfected cells and activating local bystander T_RM_; one thousand episodes simulated per model; model descriptions below the figure; episode severity measured according to **(A)** time to peak viral load, **(B)** log10 total number of infected cells by 5 days, **(C)** log10 peak viral load and **(D)** duration; results displayed with histograms showing median, interquartile range box, values within 1.5 of the IQR (whiskers) and individual episodes (dots); models with removal of single cytokine effects continue to allow rapid control of most episodes indicating that multiple, but not all antiviral functions are required to achieve rapid elimination of infection.

**Fig S4.**
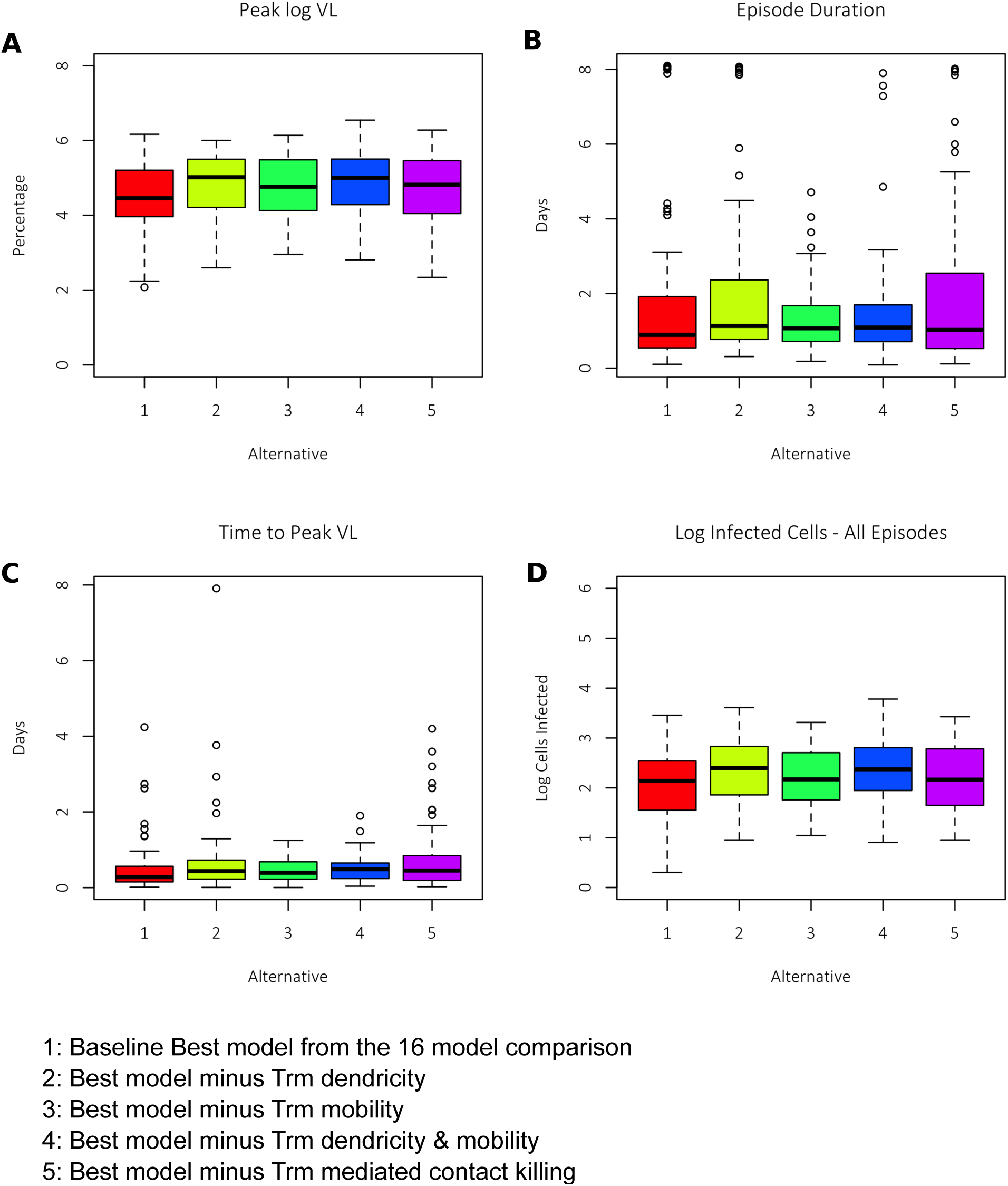
No requirement of trafficking, dendricity, and contact-mediated killing of T_RM_ for elimination of HSV-2 infected cells during severe episodes. Episodes simulated with starting E:T ratio densities randomly selected from **Fig. 2d** under conditions in which T_RM_ dendricity and / or trafficking is absent, and that T_RM_ contact-mediated killing is absent; one thousand episodes simulated per model; model descriptions below the figure; episode severity measured according to **(A)** time to peak viral load, **(B)** log10 total number of infected cells by 5 days, **(C)** log10 peak viral load and **(D)** duration; results displayed with histograms showing median, interquartile range box, values within 1.5 of the IQR (whiskers) and individual episodes (dots).

**Movie S1. Lack of containment of HSV-2 spread in the absence of T_RM_.** One simulated episode on a large cell field. Left panel: ulcer cell dynamics with cell counts in the legend, peach= uninfected cells, green = pre-productive infection, red = productive infection, black = virally lysed cell. Right panel: Viral loads (HSV DNA copies) per cellular region: purple=10^2^- 10^2.25^, blue=10^2.25^-10^2.5^, green=10^2.5^-10^2.75^, yellow=10^2.75^-10^3^, orange=10^3^-10^3.25^, red=10^3.25^- 10^3.5^, dark red>10^3.5^.

**Movie S2. Lack of containment of HSV-2 spread in the presence of T_RM_ which only kill via direct lysis of infected cells.** One simulated episode on a large cell field. Left panel: ulcer cell dynamics with cell counts in the legend, peach= uninfected cells, green = pre-productive infection, red = productive infection, black = virally lysed cell, purple=T_RM_ lysed cell. Middle panel: viral loads (HSV DNA copies) per cellular region: purple=10^2^-10^2.25^, blue=10^2.25^-10^2.5^, green=10^2.5^-10^2.75^, yellow=10^2.75^-10^3^, orange=10^3^-10^3.25^, red=10^3.25^-10^3.5^, dark red>10^3.5^. Right panel: tissue-resident T cell dynamics with cell counts in the legend, green = inactivated HSV- specific T_RM_, red = activated HSV-specific T_RM_, blue = inactivated bystander T_RM,_ purple = activated bystander T_RM_.

**Movie S3. Lack of containment of HSV-2 spread in the presence of T_RM_ which only kill via direct lysis of infected cells.** One simulated episode on a small cell field. Left panel: ulcer cell dynamics with cell counts in the legend, peach= uninfected cells, green = pre-productive infection, red = productive infection, black = virally lysed cell, purple=T_RM_ lysed cell. Middle panel: viral loads (HSV DNA copies) per cellular region: purple=10^2^-10^2.25^, blue=10^2.25^-10^2.5^, green=10^2.5^-10^2.75^, yellow=10^2.75^-10^3^, orange=10^3^-10^3.25^, red=10^3.25^-10^3.5^, dark red>10^3.5^. Right panel: tissue-resident T cell dynamics with cell counts in the legend, green = inactivated HSV- specific T_RM_, red = activated HSV-specific T_RM_, blue = inactivated bystander T_RM,_ purple = activated bystander T_RM_.

**Movie S4. Rapid containment of HSV-2 spread in the presence of T_RM_ antiviral cytokine secretion and high T_RM_ density.** Four simulated episodes with high initial T_RM_ density leading to rapid control of infection on a small cell field. Left upper panel: ulcer cell dynamics with cell counts in the legend, peach= uninfected cells, green = pre-productive infection, red = productive infection, black = virally lysed cell, purple=T_RM_ lysed cell, grey=cytokine lysed cell. Right upper panel: viral loads (HSV DNA copies) per cellular region: purple=10^2^-10^2.25^, blue=10^2.25^-10^2.5^, green=10^2.5^-10^2.75^, yellow=10^2.75^-10^3^, orange=10^3^-10^3.25^, red=10^3.25^-10^3.5^, dark red>10^3.5^. Left lower panel: tissue-resident T cell dynamics with cell counts in the legend, green = inactivated HSV-specific T_RM_, red = activated HSV-specific T_RM_, blue = inactivated bystander T_RM,_ purple = activated bystander T_RM_. Right lower panel = concentration of cytokine indicated by darkness of color; red is over infected cells or dead cells, indicating limitation of viral replication and infected cell lifespan; blue is over uninfected cells, indicating limitation of new infection.

**Movie S5. Containment of HSV-2 spread in the presence of T_RM_ antiviral cytokine secretion and low T_RM_ density.** Four simulated episodes on a small cell field with low initial T_RM_ density leading to slower control of infection. Left upper panel: ulcer cell dynamics with cell counts in the legend, peach= uninfected cells, green = pre-productive infection, red = productive infection, black = virally lysed cell, purple=T_RM_ lysed cell, grey=cytokine lysed cell. Right upper panel: viral loads (HSV DNA copies) per cellular region: purple=10^2^-10^2.25^, blue=10^2.25^-10^2.5^, green=10^2.5^-10^2.75^, yellow=10^2.75^-10^3^, orange=10^3^-10^3.25^, red=10^3.25^-10^3.5^, dark red>10^3.5^. Left lower panel: tissue-resident T cell dynamics with cell counts in the legend, green = inactivated HSV-specific T_RM_, red = activated HSV-specific T_RM_, blue = inactivated bystander T_RM,_ purple = activated bystander T_RM_. Right lower panel = concentration of cytokine indicated by darkness of color; red is over infected cells or dead cells, indicating limitation of viral replication and infected cell lifespan; blue is over uninfected cells, indicating limitation of new infection.

**Movie S6. Containment of HSV-2 spread in the presence of T_RM_ antiviral cytokine secretion and low T_RM_ density.** A simulated episode on a large cell field with low initial T_RM_ density leading to slower control of infection. Left upper panel: ulcer cell dynamics with cell counts in the legend, peach= uninfected cells, green = pre-productive infection, red = productive infection, black = virally lysed cell, purple=T_RM_ lysed cell, grey=cytokine lysed cell. Right upper panel: viral loads (HSV DNA copies) per cellular region: purple=10^2^-10^2.25^, blue=10^2.25^-10^2.5^, green=10^2.5^-10^2.75^, yellow=10^2.75^-10^3^, orange=10^3^-10^3.25^, red=10^3.25^-10^3.5^, dark red>10^3.5^. Left lower panel: tissue-resident T cell dynamics with cell counts in the legend, green = inactivated HSV-specific T_RM_, red = activated HSV-specific T_RM_, blue = inactivated bystander T_RM,_ purple = activated bystander T_RM_. Right lower panel = concentration of cytokine indicated by darkness of color; red is over infected cells or dead cells, indicating limitation of viral replication and infected cell lifespan; blue is over uninfected cells, indicating limitation of new infection.

**Movie S7. Slower containment of HSV-2 spread in the presence of T_RM_ antiviral cytokine secretion which only prevents infectivity.** Four simulated episodes. Left upper panel: ulcer cell dynamics with cell counts in the legend, peach= uninfected cells, green = pre-productive infection, red = productive infection, black = virally lysed cell, purple=T_RM_ lysed cell, grey=cytokine lysed cell. Right upper panel: viral loads (HSV DNA copies) per cellular region: purple=10^2^-10^2.25^, blue=10^2.25^-10^2.5^, green=10^2.5^-10^2.75^, yellow=10^2.75^-10^3^, orange=10^3^-10^3.25^, red=10^3.25^-10^3.5^, dark red>10^3.5^. Left lower panel: tissue-resident T cell dynamics with cell counts in the legend, green = inactivated HSV-specific T_RM_, red = activated HSV-specific T_RM_, blue = inactivated bystander T_RM,_ purple = activated bystander T_RM_. Right lower panel = concentration of cytokine indicated by darkness of color; blue is over uninfected cells, indicating limitation of new infection.

**Movie S8. Slightly protracted containment of HSV-2 spread in the presence of T_RM_ antiviral cytokine secretion which only limit lifespan of infected cells.** Four simulated episodes. Left upper panel: ulcer cell dynamics with cell counts in the legend, peach= uninfected cells, green = pre-productive infection, red = productive infection, black = virally lysed cell, purple=T_RM_ lysed cell, grey=cytokine lysed cell. Right upper panel: viral loads (HSV DNA copies) per cellular region: purple=10^2^-10^2.25^, blue=10^2.25^-10^2.5^, green=10^2.5^-10^2.75^, yellow=10^2.75^- 10^3^, orange=10^3^-10^3.25^, red=10^3.25^-10^3.5^, dark red>10^3.5^. Left lower panel: tissue-resident T cell dynamics with cell counts in the legend, green = inactivated HSV-specific T_RM_, red = activated HSV-specific T_RM_, blue = inactivated bystander T_RM,_ purple = activated bystander T_RM_. Right lower panel = concentration of cytokine indicated by darkness of color; red is over infected cells or dead cells, indicating limitation of viral replication and infected cell lifespan.

